## Supplemental Material for "Crystal Structure of a Retroviral Polyprotein: Prototype Foamy Virus Protease-Reverse Transcriptase (PR-RT)"

^1^Center for Advanced Biotechnology and Medicine (CABM), NJ, USA; ^2^Department of Chemistry and Chemical Biology, Rutgers University, Piscataway, NJ, USA; ^3^Department of Chemistry, University of Ghana, Legon, Ghana;  ^4^HIV Dynamics and Replication Program, National Cancer Institute, Frederick, MD, USA; ^5^Rega Institute and Department of Microbiology and Immunology, KU Leuven, Belgium; ^6^Department of Cell Biology and Molecular Genetics, University of Maryland College Park, College Park, MD, USA.

*Corresponding Author

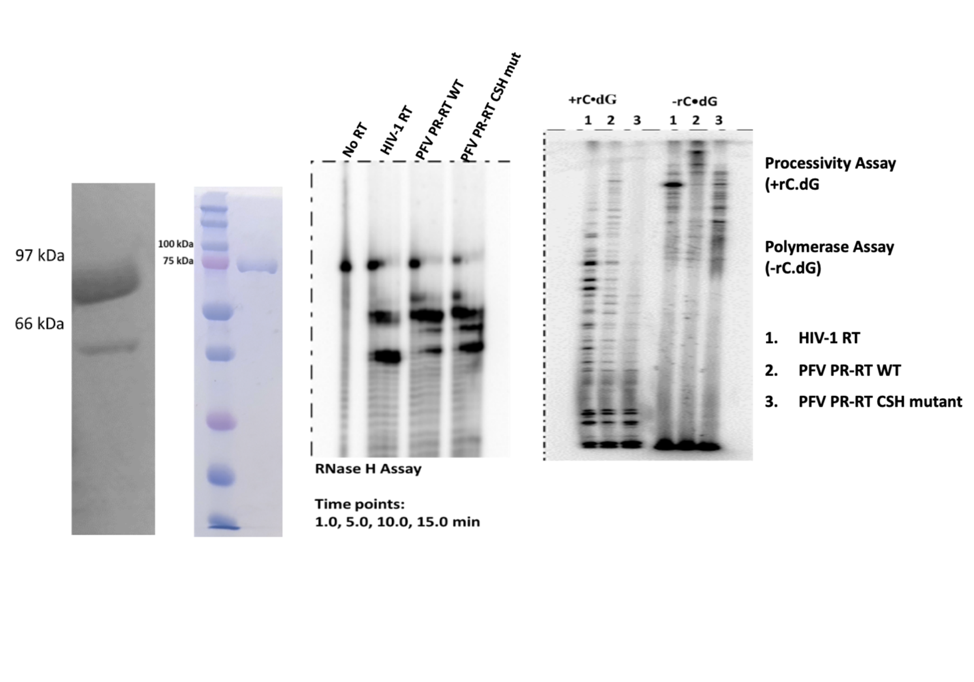

FIG S1 (A) SDS-PAGE gel of PFV PR-RT WT (left) and the CSH mutant (right) after purification. (B) Ribonuclease H activity of PR-RT and HIV-1 RT. (C) Polymerase activity and processivity (as in reference 30 of the main manuscript, Boyer et al., 2004, PMID: 15163704). The polymerase assays analyzed in three lanes on the right were done with a labeled primer annealed to single-stranded M13mp18 DNA. A polyrC/oligodG unlabeled trap which limits DNA synthesis to a single round of PR-RT binding was used. In the three lanes on the left (+rC.dG), the assays were done without a trap, and the PR-RT can rebind if it dissociates from the substrate.

C

A

B

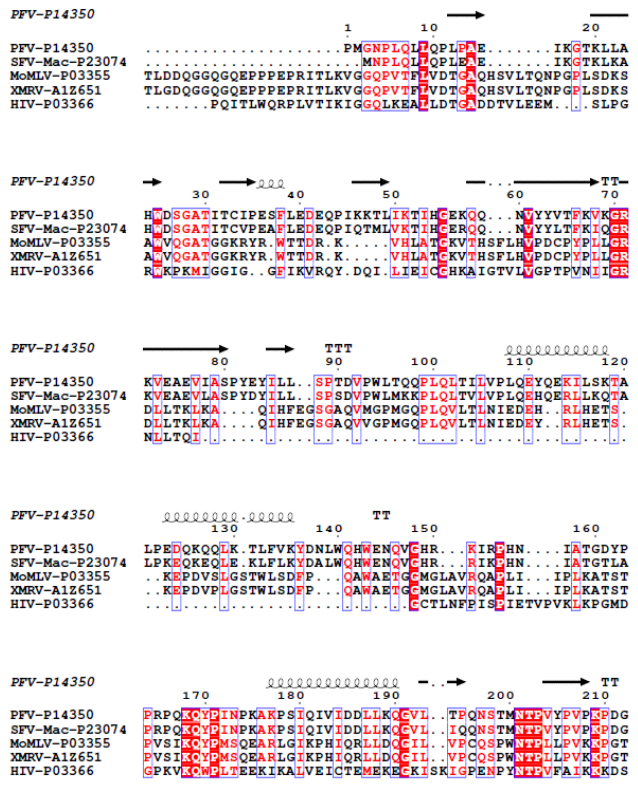

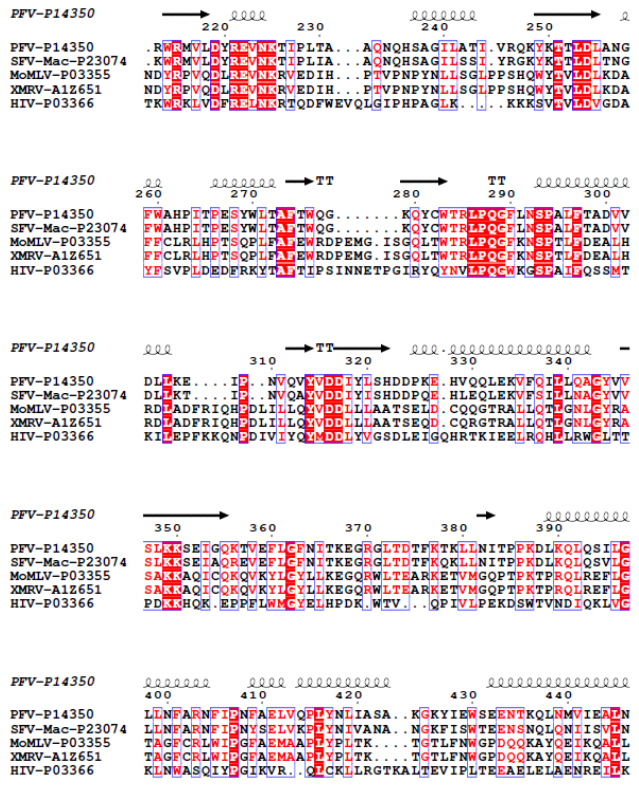

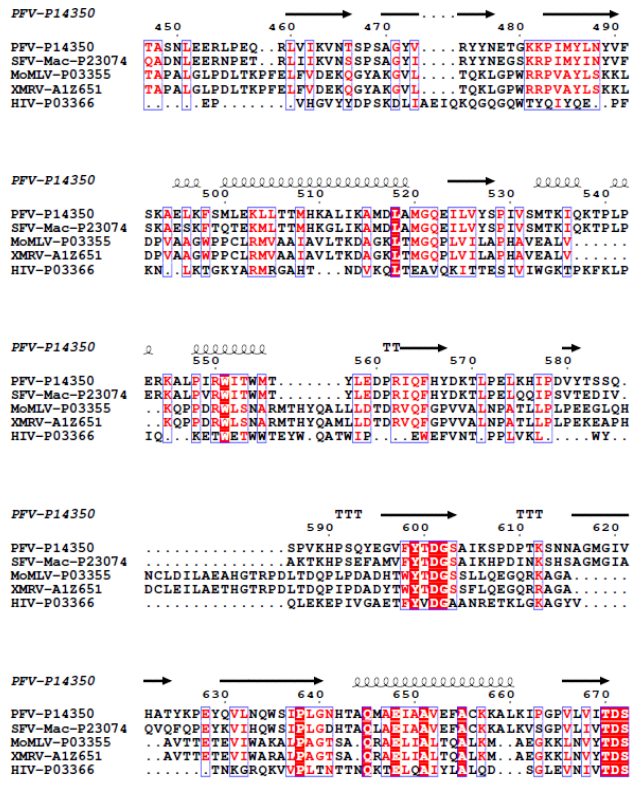

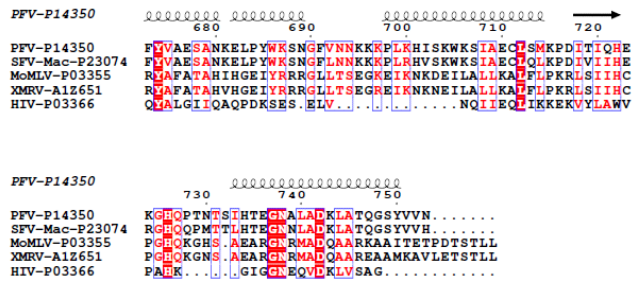

FIG S2 Alignments of a selected retroviral PRs and RTs sequences with PFV. Uniprot IDs: PFV- P14350, SFV- P23074, MoMLV- P03355, XMRV- A1Z651, and HIV-P03366. The secondary structures of PFV PR-RT and HIV-1 PR and RT are displayed at the top and bottom of aligned sequences, respectively. The primary sequences were aligned with Clustal Omega and the figure was generated using the program ESPript 3.x. Coiled lines represent α helices, solid arrows represent β strands, and Ts represent β turns.

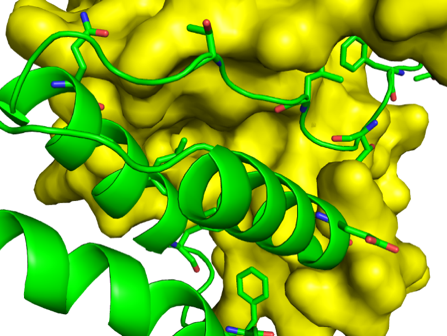

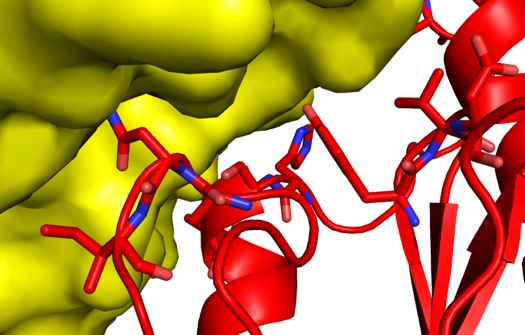

FIG S3 Left panel: interactions at the connection/thumb interface, Right panel: interactions at the connection/palm interface.

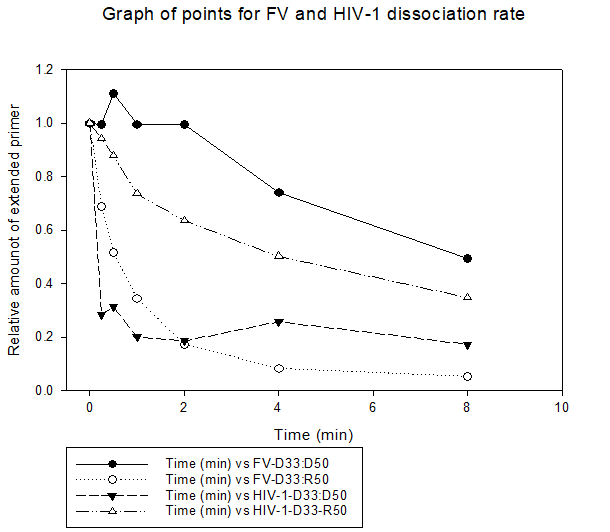

FIG S4 Dissociation rate of PFV PR-RT and HIV-1 RT from an RNA/DNA hybrid (D33-R50) and dsDNA (D33:D50). An off-rate experiment in which HIV RT or WT PFV RT was bound to a 33 nt P-32 5’ end labeled DNA primer ( 5’-TCCCCGGGTACCGAGCTCGAATTCGCCCTATAG-3’) bound to a 50 nt DNA or RNA template (same sequence: 5’DNA or RNA 5’-TTGTAATACGACTCACTATAGGGCGAATTCGAGCTCGGTACCCGGGGATC-3’) is shown. The bound RT was allowed to dissociate in the presence of a trap to prevent rebinding. Conditions used were 50 mM Tris-HCl, pH 8, 80 mM KCl, 1 mM DTT, 0.1 mM EDTA and 0.1 mg/ml BSA. The “relative amount of extended primer” on the Y-axis represents RT enzyme bound to the hybrid while time is plotted on the X-axis. For detailed methods see: Bohlayer and DeStefano, 2006 (PMID: 16768458).

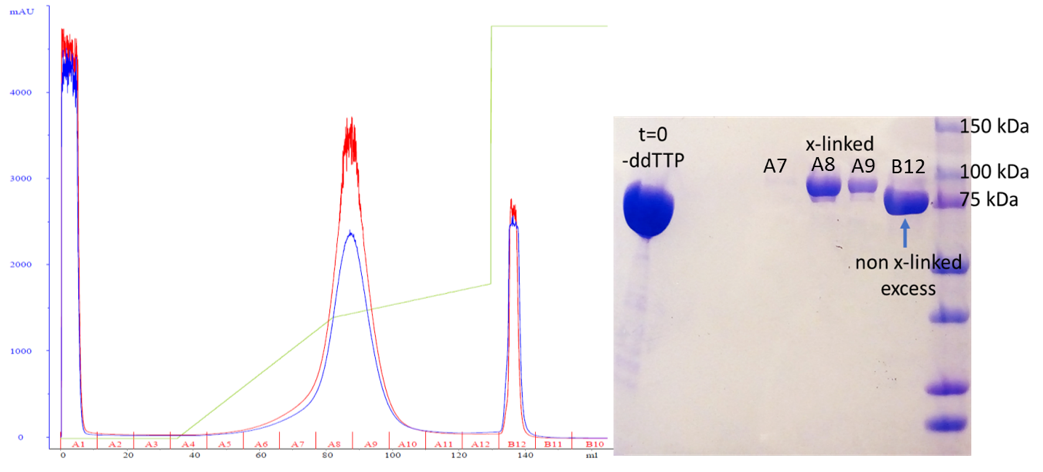

FIG S5 Heparin chromatography trace of PR-RT/dsDNA cross-linked reaction complex with its accompanying SDS-PAGE gel. Elution of the protein was done using a salt gradient.

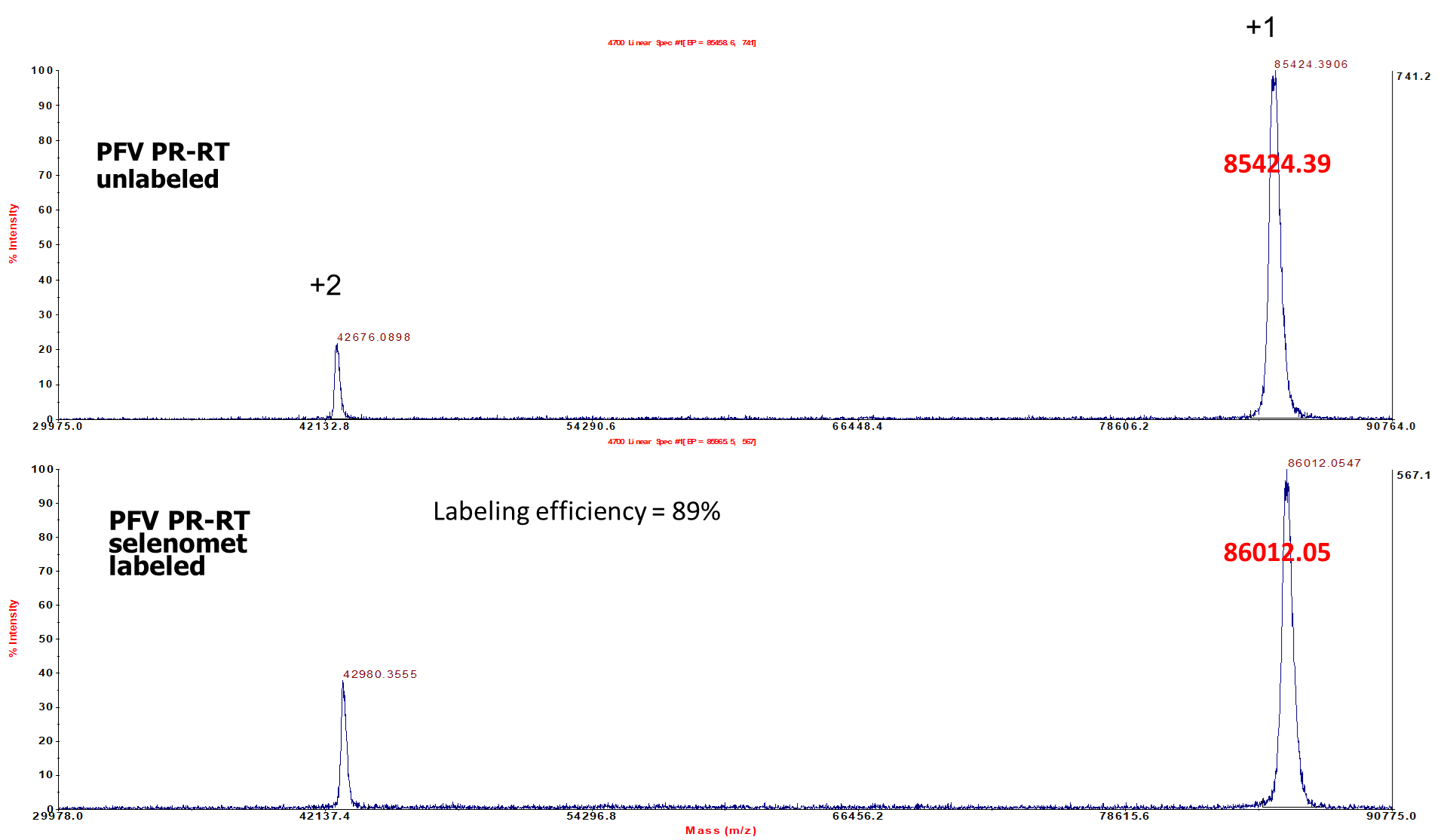

FIG S6 MALDI-TOF mass spectrum of unlabeled and SeMet-labeled PFV PR-RT.

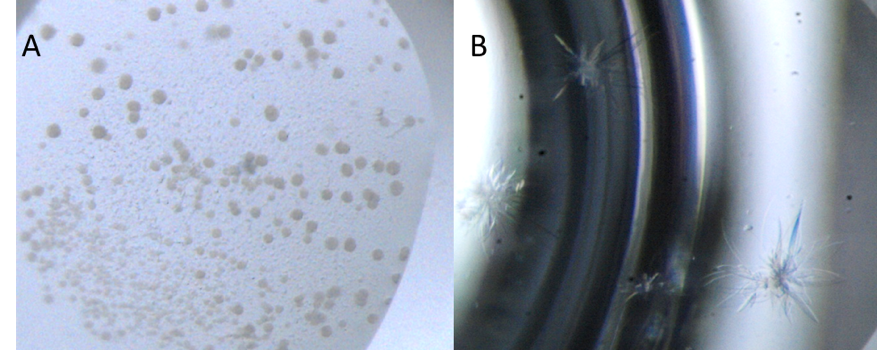

FIG S7 (A) Initial crystallization hit in Natrix HT Screen condition E8 (0.05 M potassium chloride, 0.05 M sodium cacodylate trihydrate pH 6.0, 10% w/v polyethylene glycol 8,000, 0.0005 M spermine, 0.0005 M L-argininamide dihydrochloride). (B) Optimization of initial screening Natrix HT condition E8 using 50 mM EDTA as additive.

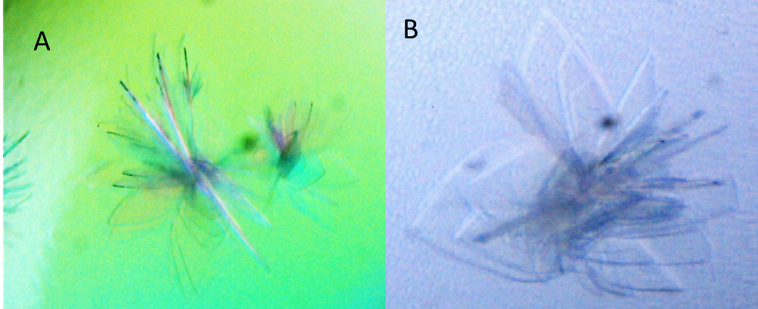

FIG S8 (A and B) Optimized crystals of PFV PR-RT used for structure determination. Optimized crystallization condition: 50 mM KCl, 50 mM sodium cacodylate trihydrate pH 6.0, 12% PEG 8000, 1.0 mM spermine, 1.0 mM L-argininamide, 200 mM glycyglycine or glycylglycylglycine, and 50 mM EDTA (or 10 mM MgCl_2_, 10 mM MnCl_2_, or 100 mM CaCl_2_).
