## Supplementary material for "Crystal Structure of a Retroviral Polyprotein: Prototype Foamy Virus Protease-Reverse Transcriptase (PR-RT)": molprobity validation of the PFV PR dimer

### Analysis output: all-atom contacts and geometry for PFV_PR_dimer_only_minimized-phenix.pdb

#### Summary statistics

| | | | | |
|---|---|---|---|---|
| All-Atom Contacts | Clashscore, all atoms: | 7.54 | | 84th percentile* (N=1784, all resolutions) |
| | Clashscore is the number of serious steric overlaps (> 0.4 Å) per 1000 atoms. | | | |
| Protein Geometry | Poor rotamers | 6 | 3.43% | Goal: <0.3% |
| | Favored rotamers | 160 | 91.43% | Goal: >98% |
| | Ramachandran outliers | 7 | 3.66% | Goal: <0.05% |
| | Ramachandran favored | 157 | 82.20% | Goal: >98% |
| | Rama distribution Z-score | -3.99 ± 0.50 | | Goal: abs(Z score) < 2 |
| | MolProbity score^ | 2.52 | | 46th percentile* (N=27675, 0Å - 99Å) |
| | Cβ deviations >0.25Å | 0 | 0.00% | Goal: 0 |
| | Bad bonds: | 0 / 1598 | 0.00% | Goal: 0% |
| | Bad angles: | 0 / 2181 | 0.00% | Goal: <0.1% |
| Peptide Omegas | Cis Prolines: | 0 / 16 | 0.00% | Expected: ≤1 per chain, or ≤5% |
| | Cis nonProlines: | 6 / 177 | 3.39% | Goal: <0.05% |
| Low-resolution Criteria | CaBLAM outliers | 14 | 7.5% | Goal: <1.0% |
| | CA Geometry outliers | 5 | 2.67% | Goal: <0.5% |
| Additional validations | Chiral volume outliers | 0/259 | | |
| | Waters with clashes | 0/0 | 0.00% | See UnDowser table for details |

In the two column results, the left column gives the raw count, right column gives the percentage.

* 100th percentile is the best among structures of comparable resolution; 0th percentile is the worst. For clashscore the comparative set of structures was selected in 2004, for MolProbity score in 2006.

^ MolProbity score combines the clashscore, rotamer, and Ramachandran evaluations into a single score, normalized to be on the same scale as X-ray resolution.

Key to table colors and cutoffs here:

##### Summary Table Cutoffs

| Category | Validation | Good | Caution | Warning |
|---|---|---|---|---|
| All-Atom Contacts | Clashscore, all atoms: | Percentile ≥ 66 | 66 > Percentile ≥ 33 | Percentile < 33 |
| Protein Geometry | Poor rotamers: | Outliers ≤ 0.3% | 0.3% < Outliers ≤ 1.5% | Outliers > 1.5% |
| | Favored rotamers: | Favored ≥ 98% | 98% > Favored ≥ 95% | Favored < 95% |
| | Ramachandran outliers: | Outliers ≤ 0.05% | 0.05% < Outliers ≤ 0.5%<br>or<br>Outliers > 0.5% and Outlier count = 1 | Outliers > 0.5%<br>and<br>Outlier count ≥ 2 |
| | Ramachandran favored: | Favored ≥ 98% | 98% > Favored ≥ 95% | Favored < 95% |
| | Ramachandran Z-score: | abs(Z-score) ≤ 2 | 2 < abs(Z-score) ≤ 3 | abs(Z-score) > 3% |
| | MolProbity score: | Percentile ≥ 66 | 66 > Percentile ≥ 33 | Percentile < 33 |
| | Cβ deviations >0.25Å: | Outlier count = 0 | 0 < Outliers < 5% | Outliers ≥ 5% |
| | Bad bonds: | Outlier bonds < 0.01% | 0.01% ≤ Outlier bonds < 0.2% | Outlier bonds ≥ 0.2% |
| | Bad angles: | Outlier angles < 0.1% | 0.1% ≤ Outlier angles < 0.5% | Outlier angles ≥ 0.5% |
| Peptide Omegas | Cis Prolines: | No suitable | universal cutoffs | for cisProline |
| | Cis nonProlines: | Peptides ≤ 0.05% Cis | 0.05% < Cis Peptides ≤ 0.1% | Peptides > 0.1% Cis |
| | Twisted Peptides: | Twisted peptide count = 0 | 0 < Twisted peptides ≤ 0.1% | Peptides > 0.1% twisted |
| Nucleic Acid Geometry | Probably wrong sugar puckers: | Outlier count = 0 | 0 < Outliers ≤ 5% | Outliers > 5% |
| | Bad backbone conformations: | Outliers ≤ 5% | 5% < Outliers ≤ 15% | Outliers > 15% |
| | Bad bonds: | Outlier bonds < 0.01% | 0.01% ≤ Outlier bonds < 0.2% | Outlier bonds ≥ 0.2% |
| | Bad angles: | Outlier angles < 0.1% | 0.1% ≤ Outlier angles < 0.5% | Outlier angles ≥ 0.5% |
| Low-resolution Criteria | CaBLAM outliers: | Outliers ≤ 1% | 1% < Outliers ≤ 5% | Outliers > 5% |
| | CA Geometry outliers: | Outliers ≤ 0.5% | 0.5% < Outliers < 1% | Outliers > 1% |

The green-to-yellow cutoffs are set from statistics or from properties of the validation methods.
E.g. the Ramachandran "Favored" category is defined by the contour containing the top 98% of residues, therefore the green-to-yellow cutoff is set at 98%.

The yellow-to-red cutoffs are not derived from properties of the validations, but are set from our intuition and experience to serve as guildlines for when structures become seriously troubled.
E.g the Ramachandran "Favored" line becomes colored red when fewer than 95% of residues fall within the 98% contour.

Ramachandran validation allows a structure with a single outlier to be colored yellow to avoid seeming to disproportionalte punish small structures.

Ramachandran Z-score validation checks the total Ramachandran *distribution* against the expected distribution. This can reveal overfitting. For more information on Ramachandran Z-score, see: https://www.biorxiv.org/content/10.1101/2020.03.26.010587v1

We determined that the expected cisProline content of strutures varies to widely for a simple cutoffs to avoid either passing or penalizing a significant number of structures incorrectly. Therefore, a structure's cisProline count is reported, but the row in the table is left uncolored.
A similar enrichment of cis nonPro is expected in certain proteins, but real cis nonPro is sufficiently rare and erroneous cis nonPro sufficiently common that we color the row to ensure that modelers are alerted to the presense of even justified cis NonPro.

As always with structure validations, "perfect" is not the correct goal. "Justified" is the goal, and outliers are not necessarily mistakes.
Especially in small structures, embrace yellow validation rows if you can explain and justify the outliers.
